# spammR: an R package designed for analysis and integration of spatial multi-omic measurements

**DOI:** 10.1101/2025.08.26.672472

**Authors:** Yannick Mahlich, Harkirat Sohi, Marija Velickovic, Paul Piehowski, Jason E McDermott, Sara J Gosline

**Author notes:** Contact* Sara Gosline.

## Abstract

**Summary:** Spatial omics is a young and evolving field and as such shows rapid development of novel technologies and analysis methods to measure transcripts, proteins, metabolites, and post-translational modifications at high spatial resolution. These advances in technology have enabled the simultaneous generation of abundance profiles for multiple different omics types and associated microscopy imaging data, as well as their analysis in a spatial context. However, most analytical tools are designed for spatial transcriptomics platforms and are challenging to use in other contexts such as mass spectrometry-based measurements or metagenomics.

To this end we present spammR (<u>sp</u>atial <u>a</u>nalysis of <u>m</u>ulti-omics <u>m</u>easurements in <u>R</u>), an R package that enables end-to-end analysis with a specific focus on mass-spectrometry derived spatial omics datasets with the goal of integration across multiple data types (e.g. sequencing, metabolites, proteins) within the same tissue sample.

**Availability and implementation:** spammR is implemented in R. The package is currently installable from GitHub (https://github.com/PNNL-CompBio/spammR).

## 1. Introduction

The field of spatial biology (Crosetto et al. 2015; Moses and Pachter 2022; Vandereyken et al. 2023) and the growth of spatially-resolved technologies (Bouwman et al. 2022; Moffitt et al. 2022; Lundberg and Borner 2019; Taylor et al. 2021) has emerged from the need to study biological systems within their spatial context and native tissue environments (Kiessling and Kuppe 2024). Measuring the molecular changes of a sample, e.g. slices of tumor tissue, in its spatial context (Nichterwitz et al. 2018; Lee et al. 2014; Zhang et al. 2021), has been enabled by both novel transcriptomics (Eng et al. 2019; Rodriques et al. 2019; Codeluppi et al. 2018; Chen et al. 2015; Wang et al. 2018) as well as proteomics technologies (Radtke et al. 2022; Black et al. 2021; Angelo et al. 2014; Lin et al. 2018). Mass spectrometry (MS) based technologies to measure proteins (Zhu et al. 2018; Piehowski et al. 2020), lipids, metabolites, and other molecules (Buchberger et al. 2018; Greer et al. 2011; Veli_č_kovi_ć_ et al. 2024; 2025) have furthered enabled this field to study how molecular interactions can vary across a tissue.

While the pace of computational tool development has accelerated, particularly in the field of spatial transcriptomics (Walker et al. 2022), there is a gap in spatial analysis tools tailored towards the analysis and interpretation of mass spectrometry-based multi-omics data, with most existing tools focused on specific analytical frameworks (Nimo et al. 2025) that cannot be readily applied to integrative omics analyses – e.g. comparing metabolites and proteomics in the same sample. This is especially important regarding the distinct differences between antibody imaging and mass spectrometry-based measurement. For mass spectrometry specifically, those include (1) the need for normalization of mass/charge ratios, (2) the potential need to impute data to counteract missingness and, (3) MS as a technology not being restricted to proteomics. In addition to MS-based spatial proteomics, the rise of spatial glycomics, metabolomics, lipidomics and measurements of post translational modifications (PTMs), such as phosphorylation, all necessitate a more flexible framework than those that exist for other spatial technologies.

To this end, we introduce spammR (SPatial Analysis of Multiomics Measurements in R), an R package that is well-suited for an end-to-end analysis of mass-spectrometry derived spatial omics datasets with (1) smaller sample sizes and spatial sparsity of sampling, (2) considerable missingness, and (3) no a-priori knowledge about proteins or genes of interest, relying on a fully data-driven approach. Here we describe the overall features of spammR, and how it can be used in “traditional” spatial omics contexts, using proteomics data from pancreatic cancer, and lipidomics and proteomics data from rat brain as examples, as well as more broadly as a tool to perform comparative omics studies in a spatial context using metagenomic data from the 1000 soils project (Bowman et al. 2023; Song et al. 2025).

### 2. Implementation

spammR is a package implemented in R and will be available via Bioconductor. We make use of the ‘SpatialExperiment’ (Righelli et al. 2022) object via Bioconductor (Gentleman et al. 2004; Huber et al. 2015) to provide the ability to integrate with other tools in the Bioconductor environment, and for increased interoperability with other tools in the spatial omics field (Feng et al. 2023; Pardo et al. 2022; Windhager et al. 2023). The package consists of five main components (**Figure 1A**): (1) Data ingest, (2) data processing, (3) feature selection, (4) functional enrichment, and (5) integrated visualization.

**Figure 1.**
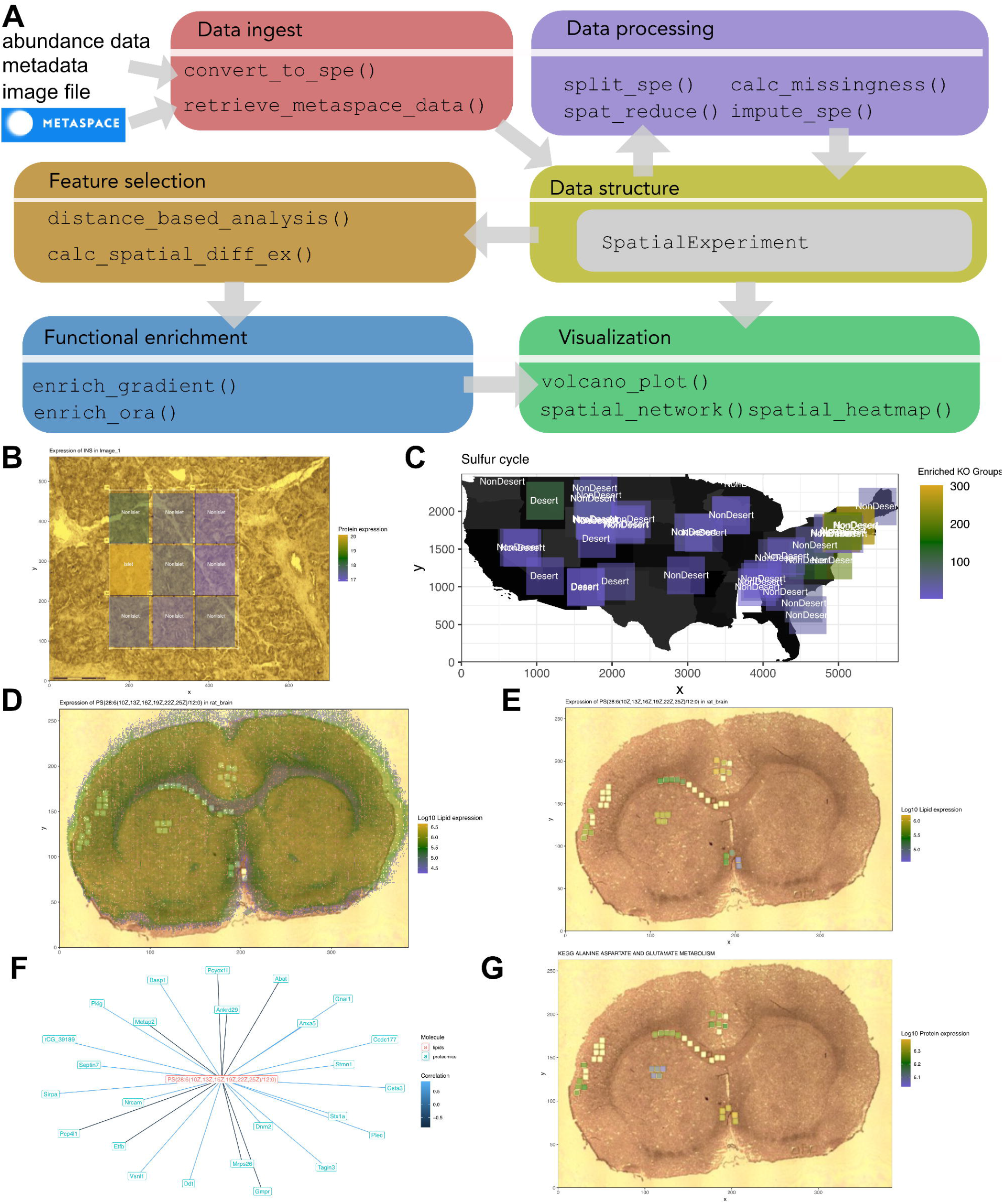
Overview of spammR functionality. (A) Flow chart describing use of spammR package as well as how the methods of the package relate to one another. (B) Expression of the protein insulin in a human pancreas tissue, with labels indicating whether the image region contains islet cells or not. (C) Abundance of the sulfur cycle KEGG orthology group across the US Soil study. (D) Expression of a highly variable lipid across a rat brain sample, as well (E) the expression of that lipid only within the regions measured by proteomics. (F) Network representing the lipid of interest (pink) and highly positively or negatively correlated proteins (turquoise). (G) Expression of proteins in the KEGG Alanine, aspartate, and glutamate pathway, which was identified as significantly enriched in proteins negatively correlated with the lipid of interest shown in panel.

The data ingest component is comprised of two functions designed to capture the diverse data modalities that spammR analyzes. The primary function is `convert_to_spe` ()and consumes three types of data: a data matrix of counts or values representing the samples measured (samples as columns, features as rows), a metadata table mapping the samples to a coordinate system as well as any other phenotypic data, and the image itself. spammR assumes that the molecular data are fully normalized/batch corrected (Välikangas et al. 2018), however additional corrections can also be applied depending on the data modality consumed (e.g. RNASeq vs proteomics). The function then returns a `SpatialExperiment` object containing the information that can be used for additional processing or analysis. To account for mass spectrometry imaging data on the Metaspace2020 platform(Veli_č_kovi_ć_ et al. 2021), we also included a `retrieve_from_metaspace` () function that selects a specific project and downloads both the coordinate metadata as well as the molecular quantifications. This function also returns a `SpatialExperiment` object. Each function only retrieves one data modality at a time, with merging happening using utility functions described below.

The data processing component currently consists of four functions designed to aid in multiomic analysis. To account for missingness in data, we included the `calc_missingness` () function to identify missing feature values in the object, and the `impute_spe()` that can run eight basic imputation algorithms. Another key feature of the spammR is the ability to measure diverse data modalities and samples. For this we included the `spat_reduce` () function that merges two SpatialExperiment objects on the same coordinate space as well as the `split_spe` () function that separates out a single object into multiple smaller objects.

The feature selection component contains functions to identify features (e.g. proteins) that either show strong correlation between abundances and spatial distance based on ROIs (`distance_based_analysis` ()) or display significant differential expression profiles based on sample categorization (e.g. a categorical distinction between ROIs) without incorporation of distance measures `(calc_spatial_diff_ex()`). Both functions take a `SpatialExperiment` object created by `convert_to_spe` () and return an augmented `SpatialExperiment` with the added significance rankings.

The functional enrichment component contains functions to generate over-representation statistics (ORA) for feature selection results. `enrich_gradient` () can be used to generate pathway enrichment based on results from the distance-based analysis (`distance_based_analysis` ()). Analogously, `enrich_ora` () does the same for ranking results from categorical differential expression analysis. Both functions rely on leapR (Danna et al. 2021) for pathway and gene set enrichment analysis (GSEA) (Subramanian et al. 2005). Like the feature selection functions, both functions in the enrichment component will return an augmented `SpatialExperiment` object that can in turn be used for visualization of enriched pathways in ROIs.

Finally, the data contained in a `SpatialExperiment` object, either directly generated by the raw data ingestion or augmented using the feature selection components, can then be visualized using the visualization component which contains two different visualization functions. (1) `spatial_heatmap` () overlays a grid onto image data representing the different ROIs and shading the individual grid cells by the calculated differential (e.g. raw abundance measures or differential expression calculated by one of the functional enrichment components). (2) `volcano_plot` () can be used to visualize the differential expression results generated by `calc_spatial_diff_ex` (). We also enable network-style visualizations through the `spatial_network` ()function, which identifies pair-wise relationships between molecules across space that can then be plotted using the tidygraph R package.

All functions are designed to be lightweight, data agnostic, and provide interoperability via the `SpatialExperiment` object to enable usage beyond the functionality outlined here.

### 3. Application

We included in the spammR package three examples of applications using multiomics measurements across space.

#### 3.1. Coordinate-based spatial heatmap in spammR enables contextual interpretation of spatial differences

To demonstrate the capabilities of spammR we conducted an analysis on previously published multi-omics data of human pancreatic tissue samples (Gosline et al. 2023). Specifically, we start by using the available methods in the spammR package to generate and visualize abundance differences for insulin across regions of interest (ROIs) which predominantly contain islets or non-islet cells respectively, observing the expected behavior of higher abundance of insulin in islet grid cells (**Figure 1B**). Next, we performed a pathway enrichment analysis using spammR’s feature selection and functional enrichment components by pooling images from 7 distinct pancreas samples. As expected, pathways associated with insulin regulation and secretion are enriched islet cells in comparison to pancreatic regions containing predominantly non-islet cells. Finally, using the distance-based analysis functions in spammR we were able to identify and visualize the fact that AP3S2, a subunit of the AP-3 complex that is thought to be associated with protein transport (Cowles et al. 1997), expression strength is strongly anti-correlated with the distance from islet cells, i.e. decreasing expression with increasing distance. The entire analysis is showcased in one of the vignettes (spatProt.rmd) included with the spammR package.

### 3.2 spammR’s agnostic approach to multi-omics integration enables the investigation of metagenomic data in a geospatial context

The implementation of spammR’s spatial component is inherently agnostic to whether the distances within the investigated sample are on a micrometer, millimeter or even kilometer scale. To demonstrate this, we retrieved KEGG (Kanehisa et al. 2025) Ortholog (KO) presence data for 108 metagenome samples (54 sampling sites, samples each at two depths) from the 1000 soils project pilot study of MONet (Molecular Observation Network). Note that this is different from mass-spectrometry proteomics data in so far that in the 1000 soils data the number of predicted protein sequences from metagenome-assembled genomes (MAGs) that are associated with a given KO is recorded (i.e. presence), whereas traditional MS-proteomics data reports quantitative abundance measurements. This leads to significantly lower “abundance values” as well as lower variance between the individual values. In addition to the KO abundance, each sample contains further metadata, like soil temperature, water content, pH, metal abundances, and most importantly for spatial context, latitude, longitude, elevation and whether the sample originates from the top or the bottom of the extracted soil core. Using the metadata, we retrieved 575 KOs that show significant differential abundances across samples from desert vs. non-desert sampling locations (483 “up regulated”, 92 “down regulated”). We then utilized leapR’s functionality to retrieve enriched KEGG Pathways based on the KO abundance data. The pathway enrichment detected several enriched pathways, for example “sulfur cycle”. Utilizing map data from the US census bureau, the terra R package as well as the latitude and longitude annotations we are then able to visualize the pathway enrichment using the `spatial_heatmap` () function (**Figure 1C**). A complete workflow can be found in the spatMicrobiome vignette that is part of spammR.

### 3.4 Integration of proteomic and lipidomic measurements in brain tissue

To show how spammR can be used to integrate multiple omics from the same sample, we leveraged data from Vandergrift et al. (Vandergrift et al. 2025) that measured lipidomics and proteomics from a tissue sample from rat brain. We used the `retrieve_metaspace_data` () to fetch the experimental data from the Metaspace portal (Palmer et al. 2017) and coordinates from the lipids, then downloaded the proteomics data from the original publication. The original lipid data, shown in **Figure 1D**, spans the entire image, while the protein data (carved out as squares in the image) only represented a subset of the image. We therefore used the spat_reduce() function to align the two coordinate systems to map the average lipid expression to the regions that correspond to protein measurements, shown in **Figure 1E**. Using this combined object, we could then build a correlation graph, shown in **Figure 1F**, that identifies specific proteins correlated with a particular lipid of interest, in this case a phosphatidylserine identified as highly variable across the image (PS(28:6(10Z,13Z,16Z,19Z,22Z,25Z)/12:0)). We then ran leapR analysis on the correlation results to identify a KEGG pathway significantly enriched (negatively correlated in this case) with the lipid, depicted in **Figure 1G**.

## Conclusion

Through spammR, we address multiple limitations in the current state-of-the-art tools for computational analysis of diverse spatial omics measurements with a suite of lightweight methods that can be broadly applied across mass spectrometry and sequence-derived measurements. spammR is built upon the existing Bioconductor framework for spatial omics measurements and can therefor interoperate with the ever-growing set of tool for spatial transcriptomics, but adds specific expansions to explicitly (a) account for issues that generally arise with mass spectrometry such as missingness and (b) align omics from diverse modalities in the same image.

We worked to include a diverse set of features that could be used in the spatial omics community, but made distinct choices about when to include functionality vs. provide an interface to others. For example, in the design of our `impute_spe` () we tried to address the importance of missing data (e.g. in data-dependent acquisition (DDA)(Wu and MacCoss 2002) proteomics data). In general, missing abundance for individual features does not equate to proteins not being expressed and can broadly be categorized into values missing (completely) at random (MCAR & MAR) and values missing not at random (MNAR). MAR features can mostly be attributed to experimental design or instrument errors. Hence, missing value intensities do not necessarily follow a pattern, making them more difficult to impute. On the other hand, MNAR values can often be traced back to technical limitations of the instrument, e.g. proteins of low expression not being detected due to intensities below the detection capabilities of the instrument. Given the different nature of MAR and MNAR values where missing values are likely to follow different intensity distributions, existing imputation strategies do not universally perform well for both types of missing values(Lazar et al. 2016; Webb-Robertson et al. 2015; Wei et al. 2018). For example, assuming MNAR for observed missing values, naïve single value imputation methods like SampMin (minimal value of the sample) can show good performance especially in downstream analyses like differential expression analysis(Harris et al. 2023; Kong et al. 2022; Liu and Dongre 2021). Especially when dealing with spatial data, where cell type-specific proteins that may only be expressed in one region of the tissue, those naïve imputation strategies are a possible approach. To account for diverse use cases, `impute_spe` () currently includes eight imputation methods including spatially-aware imputation method that adapts k-nearest neighbor imputation to assign values to missing abundance data points based on molecular abundances only from samples that are spatially close to each other (rather than close to each other in expression value). Finally, considering that spammR is built on widely used data structures with a focus on interoperability with other R packages, users also have the ability to integrate other available imputation strategies from libraries like imputeLCMD(Lazar et al. 2022) or MsCoreUtils(Rainer et al. 2022) into their workflow.

We demonstrated the power of spammR for the analysis of a traditional spatial proteomics datasets using a spatial proteomics analysis of healthy human pancreatic tissue. Additionally, we showed that spammR is scale agnostic and can also be utilized on geospatial distance scale data by using a metagenome dataset highlighting the agnostic capabilities of spammR extending beyond the analysis of traditional multi-omic data. Lastly, we showed the power of integration using a small dataset that collects both lipidomics and proteomics from the same tissue.

In the future, we plan to continue expanding the functionality provided by spammR through the collection of larger, more complex datasets across tissues of interest. We aim to improve data processing through the incorporation of other spatial analysis such as the `SpatialFeatureExperiment` (Moses et al. 2023) and the direct incorporation of geoJSON format to link to tools such as quPath (Bankhead et al. 2017) or microscope vendor software. As multiomics datasets become more comprehensive, we also plan to improve our statistical methods for identification of functional pathways that account for measurements across omics.

## Acknowledgements

Soil data were provided by the Molecular Observation Network (MONet) at the Environmental Molecular Sciences Laboratory (https://ror.org/04rc0xn13), a DOE Office of Science user facility sponsored by the Biological and Environmental Research program under Contract No. DE-AC05-76RL01830. The work (proposal: 10.46936/10.25585/60008970) conducted by the U.S. Department of Energy, Joint Genome Institute (https://ror.org/04xm1d337), a DOE Office of Science user facility, is supported by the Office of Science of the U.S. Department of Energy operated under Contract No. DE-AC02-05CH11231. A subset of soil cores was obtained by the National Ecological Observatory Network’s Research Support Services which is a program sponsored by the U.S. National Science Foundation and operated under cooperative agreement by Battelle. The Molecular Observation Network (MONet) database is an open, FAIR, and publicly available compilation of the molecular and microstructural properties of soil. Data in the MONet open science database can be found at https://sc-data.emsl.pnnl.gov/.

## Author contributions

**Yannick Mahlich:** Analysis, Writing – original draft, **Harkirat Sohi:** Methodology, Software, **Jason McDermott:** Conceptualization, Supervision, Writing – editing, **Sara Gosline:** Conceptualization, Methodology, Supervision, Software, Writing – original draft

## Funding

This work was supported by DoD CDMRP Program W81XWH-22-1-0324, NF210042; NIH grant U01-CA271412; and USDA NF-SPP 81327.

## Conflict of interest

None

## Notes

https://github.com/pnnl-CompBio/spammR

### Competing Interest Statement

The authors have declared no competing interest.

### Summary of Updates

An additional analysis was provided as an example and figure was updated.

