## Supplemental Materials for "spammR: an R package designed for analysis and integration of spatial multi-omic measurements"

**Supplementary Methods**

*Dataset descriptions*

We chose to include three published datasets for the purpose of demonstration of this R package:

1. Pancreas proteomics: This published dataset (Gosline et al. 2023) is comprised of seven images, each with nine regions (voxels) arranged in a 3x3 grid format. In each image, one of the nine voxels is centered around a cluster of islet cells. The coordinates of each voxel were calculated manually and saved in an accompanying table that was used as sample metadata in the construction of the SpatialExperiment object alongside the image and a counts matrix for the normalized protein measurements for that image.
2. Global microbial soils measurements: This published dataset(Bowman et al. 2023) was retrieved from the Molecular Observation Network (MONet) pilot project (1000 soils). KEGG Ortholog abundances for the 1000 soils project data were provided by Song et al.(Song et al. 2025). Geospatial coordinates as well as any metadata for all of the soil core microbiomes (e.g. sampling depth, site biome, moisture content, etc.) was retrieved using the MONet visualization tool’s ‘QUERY’ and ‘SITE_INFORMATION’ tabs (<https://shinyproxy.emsl.pnnl.gov/app/monet-vis-testbed>). Map data was retrieved from the US census bureau (<https://www.census.gov/cgi-bin/geo/shapefiles/index.php>). Coordinates were mapped using the terra and sf R packages. KO abundance data, metadata tables and the generated map image of the continental US were used in the construction of a SpatialExperiment object. A more detailed description of the process can be found in spammR’s spatMicrobiome vignette.
3. Lipidomics and proteomics from rat brain: This published dataset(Vandergrift et al. 2025) includes MALDI-based lipidomic measurements together with proteomics measurements from several regions of interest (demarcated by squares in the image file). The image file was downloaded manually from Metaspace2020 together with the lipid measurements which were downloaded using the retrieve_metaspace_data() function (dataset PNNL05A_V6b_CLMCAFAMM_Lipids_885_1ppm). The image file was imported into QuPath(Bankhead et al. 2017) to select the ROIs, which were exported into GeoJSON format and imported into spammR in the vignette. We then merged the two datasets using the spat_reduce() function to carry out enrichment analysis.
